# Atypical cortical encoding of the low-frequency temporal dynamics of natural speech identifies children with Developmental Language Disorder

**DOI:** 10.64898/2026.03.08.710292

**Authors:** Xuanci Zheng, João Araújo, Mahmoud Keshavarzi, Georgia Feltham, Susan Richards, Lyla Parvez, Usha Goswami

**Author notes:** **Correspondence:** Xuanci Zheng.

## Abstract

Developmental Language Disorder (DLD) is a neurodevelopmental condition that causes significant difficulties in understanding and using spoken language. Here we use electroencephalography (EEG) recorded during natural speech listening with 9-year-old children to identify dynamic neural processing patterns that characterize children with DLD compared to typically-developing age-matched controls. We applied Principal Components Analysis (PCA) to identify spatial ensembles of channels that represented distinct (uncorrelated) sources of cortical activity, and explored phase-amplitude coupling (PAC) between delta (0.5-4 Hz), theta (4-8 Hz) and low-gamma (25-40 Hz) oscillations. We then isolated EEG common spatial patterns (CSP) that identified children with DLD. The PAC analyses identified the delta band as a key source of group differences, and only delta-low gamma PAC differed significantly between participant groups. The CSP analyses also identified the delta band as a key mechanistic source of group differences. The findings are suggestive of distinct atypical low-frequency neural dynamics during speech encoding for children with DLD, which could be targeted by novel interventions.

---

Children diagnosed with developmental language disorder (DLD) present with language difficulties that cause functional impairments in many aspects of linguistic processing (Leonard, 2014; Bishop, 2013). For example, the CATALISE multi-national study of DLD identified linguistic processing problems with syntax and grammar, a limited understanding of word meanings, impairments in verbal short-term memory, and difficulties with phonology (word sound structure, Bishop et al., 2017). Given this wide range of impairments, the underlying biomedical aetiology seems likely to involve fundamental sensory/neural aspects of speech processing. The current study follows Araújo et al. (2024) in using Temporal Sampling (TS) theory (Goswami, 2011) and electroencephalography (EEG) recorded during natural speech listening to identify neural processing patterns during connected speech listening that may identify children with DLD. TS theory is based in part on atypical features of sensory and linguistic processing found in children with DLD, including decreased sensitivity to amplitude envelope rise times (AERTs, see Corriveau et al., 2007; Corriveau & Goswami, 2009; Fraser et al., 2010; Beattie & Manis, 2012), which automatically trigger neural encoding of the speech signal (see Giraud & Poeppel, 2012). Linguistically, children with DLD show decreased sensitivity to syllable stress patterns and decreased sensitivity to speech and non-speech rhythm patterns, associated with impaired AERT discrimination (Richards & Goswami, 2015, 2019; Cumming et al., 2015a,b). The TS framework adopts a neural oscillatory speech processing framework drawn from the adult literature to explain these atypical features, and predicts impaired neural dynamics in DLD focused on the low-frequency oscillations <10 Hz that are critical for encoding speech rhythm information (EEG delta and theta bands).

Low-frequency oscillations appear to be critical for the initial set-up of an efficient speech processing system by infants (Goswami, 2022), and hence are a plausible source of inefficient speech processing in DLD. Ever since sensitivity to speech rhythm was identified as a cognitive cross-language precursor of language acquisition (Mehler et al., 1988), behavioural investigations have shown a central role for rhythm in linguistic development. For example, infants can distinguish languages with different speech rhythm patterns from birth (e.g., French versus Dutch, see Nazzi et al., 1998), they babble using the rhythm patterns of the ambient language (e.g., French versus Arabic, see Boysson-Bardies et al., 1984), and prosodic templates (the dominant rhythmic patterns that denote words in a language, such as the strong-weak template that characterizes nouns in English and German) appear to be encoded as early as 4 months of age (Weber et al., 2004). Recently, the neural oscillatory speech processing framework originally developed on the basis of adult MEG and EEG studies (Ahissar et al., 2001; Poeppel, 2003; Giraud & Poeppel, 2012; Gross et al., 2013) has been applied to infants. This framework identified the neural oscillatory bands that track different temporal features of human speech in adult cortex as primarily delta (0.5-4 Hz), theta (4-8 Hz), beta (15-25 Hz) and low gamma (30Hz+), thought to relate to prosodic information, syllabic information, speech-motor information and phonemic information respectively. These different linguistic features of speech relate to temporal modulations at different rates which are nested in the speech amplitude envelope (AE), the slow-varying energy profile of the acoustic waveform (Houtgast & Steeneken, 1985). Key roles were proposed for the theta and gamma bands regarding adult speech encoding (e.g., Giraud & Poeppel, 2012), as adult studies focused on syllabic and phonemic parsing, with theta-gamma phase-amplitude coupling (PAC) proposed as an important mechanism for integrating these rhythms during efficient speech perception (e.g., Dogonasheva et al., 2025 for recent review).

Insights from adult auditory neuroscience are beginning to be applied to language acquisition. The infant brain tracks the speech AE from birth onwards (Kalashnikova et al., 2018; Jessen et al., 2019; Ortiz Barajas et al., 2021; Attaheri et al., 2022; Menn et al., 2022). However, while the AE of adult speech in different languages shows a modulation peak at ∼5 Hz in the theta band across languages (Varnet et al., 2017, Ding et al., 2017), infant-directed speech shows a modulation peak at ∼2Hz in the delta band (Leong et al., 2017). This suggests that oscillatory delta-band prosodic-level information is unconsciously foregrounded when speaking to infants, and hence may be critical for the initial formation of a linguistic system. In a recent longitudinal study of 120 infants followed from the age of 2 months, the accuracy of delta band encoding of connected speech (nursery rhymes) was significantly greater than the accuracy of theta band encoding at 4, 7 and 11 months. Accuracy of delta band encoding at 11 months predicted language outcomes at age 2 years, but the accuracy of theta band encoding did not predict language outcomes (Attaheri et al., 2024). A second predictor of language outcomes at age 2 years was theta-gamma PAC at age 4 months (Attaheri et al., 2024). Accordingly, the accuracy of both delta band encoding and the efficiency of PAC between slower and faster neural rhythms are plausible targets for exploring the neural mechanisms that may be affected in DLD.

The analyses reported here were modelled on a classifier study of 16 children with developmental dyslexia reported by Araújo et al. (2024). In that study two groups of children acted as control groups, a typically-developing (TD, N=40) group and a group with DLD (N=7). Araújo et al. (2024) used a combined approach of principal component analysis (PCA) and common spatial patterns (CSP) to analyse EEG data recorded during a passive story listening task. This initial study found that the DLD group showed lower delta-theta PAC variance (*p*= .022, corrected) compared to the TD controls for the first of the three dominant principal components, which related to frontal activation. The DLD group also showed a significantly higher mean delta-theta PAC value for the second principal component relating to bilateral temporal activation (*p*= .002, corrected) when compared to the TD controls. These data suggested that low-frequency PAC metrics may distinguish children with DLD from TD controls during continuous speech listening. When CSPs were examined, TD children exhibited greater EEG power in the delta and theta bands than DLD children in central and left-lateralized regions, while a spatial filter focusing on occipital channels maximzed the variance for children with DLD in the *beta band* range. These data were interpreted to suggest that atypical delta-theta cross-frequency coupling may identify children with DLD in natural speech listening tasks, along with atypical beta band activity in occipital regions. However, as only 7 children with DLD provided data for this study, replication with a larger sample is required.

In the current study, we examine whether atypical delta-theta PAC and occipital CSP features in the beta band will distinguish children with DLD from TD controls. We employ a larger sample of children with DLD compared to Araújo et al. (2024) and we use a different story in the story listening task while recording EEG (Keshavarzi et al., 2022). We also use identical PAC methodology to Araújo et al. (2024), who used a method that does not rely on bandpass filtering. Accordingly, if similar atypical neural dynamics are found in the current study as in Araújo et al. (2024), this could be suggestive of fundamental oscillatory differences in the DLD vs TD brain during active speech processing. We here (a) derive group-blind spatial filters for the EEG data using PCA, (b) compute phase-amplitude coupling for each principal component between delta-theta, delta-low gamma and theta-low gamma to explore how low-frequency phases couple with both low-frequency (theta) and high-frequency (gamma) power, and (c) derive CSPs that maximize the variance for the TD group and minimize the variance for the DLD group and vice versa, here analysing all EEG bands (delta, theta, alpha, beta, low gamma). We then aim to classify the children with DLD versus TD control children on the basis of our findings. On the basis of TS theory, *a priori* we expect that delta-driven oscillatory dynamics may play a key role in the aetiology of DLD.

## 2. Materials and methods

### 2.1. Participants

Sixteen typically developing children (TD control group, mean age of 9.1 ± 1.1 years) and sixteen children with DLD (mean age of 9 ± 0.9 years) contributed data for the study. All participants were taking part in an ongoing study of auditory processing in DLD (Parvez et al., 2024), and those children with suspected language difficulties were nominated by the special educational needs teachers in their schools. Children in the TD group were nominated by classroom teachers as being typically-developing. All children had English as the main language spoken at home. All participants exhibited normal hearing when tested with an audiometer. In a short hearing test across the frequency range 0.25 – 8 kHz (0.25, 0.5, 1, 2, 4, 8 kHz), all children were sensitive to sounds within the 20 dB HL range. Language status was ascertained by adminstering two subtests of CELF-V (Wiig, Semel & Secord, 2017) to all the participants: recalling sentences and formulating sentences. Those children who appeared to have language difficulties then received (depending on age) two further CELF subtests drawn from word structure, sentence comprehension, word classes and semantic relationships. Children who scored at least 1 S.D. below the mean (7 or less when the mean score is 10) on at least 2 of these 4 subtests were included in the DLD group. Oral language skills in the control children were thus measured using only two CELF tasks, recalling sentences and formulating sentences, and all TD control children scored in the normal range (achieving scores of 8 or above). Due to the Pandemic, although all the control children received the recalling sentences task, only 11 also received the formulating sentences task (all children with DLD received 4 CELF tasks). The Picture Completion subtest of the Wechsler Intelligence Scale for Children (WISC IV, Wechsler, 2016) was used to assess non-verbal intelligence (NVIQ) and did not differ significantly between groups. Group performance for the tasks administered to both groups is shown in Table 1. All participants and their parents provided informed consent to participate in the EEG study, in accordance with the Declaration of Helsinki. The study was reviewed by the University of Cambridge, Psychology Research Ethics Committee and received a favourable opinion.

**Table 1.** Details of the participating children, showing Scaled Scores (ScS, population mean = 10).

|  | DLD | TD Controls | Significance |
| --- | --- | --- | --- |
| N | 16 | 16 | — |
| Age (months) | 108.3 (11.3) | 108.9 (13.3) | 0.585 |
| Recalling Sentences ScS | 5.3 (1.7) | 10.9 (2.3) | 0.0001 |
| Formulating Sentences ScS | 4.4 (1.9) | 10.8 (1.7) | 0.0001 |
| WISC Picture Completion ScS | 7.9 (3.0) | 9.9 (2.8) | 0.075 |

Note. WISC: Wechsler Intelligence Scale for Children.

### 2.2. Experimental set up and stimuli

The experimental setup and stimuli utilised in the EEG study were identical to those described by Keshavarzi et al. (2022). The participants listened to a 10-minute story for children, *The Iron Man: A Children’s Story in Five Nights* by Ted Hughes. During the experiment, participants were instructed to listen to the speech carefully and to look at a red cross (+) shown on the screen that was in front of them. The sampling rate was 44.1 kHz for presenting the auditory stimulus, and EEG data were concurrently collected at a sampling rate of 1 kHz using a 128-channel EEG system (HydroCel Geodesic Sensor Net) with manufacturer-defined electrode labels (E1–E128). To ensure the children attended to the task, they were informed beforehand that they would be asked two simple comprehension questions after the story. Comprehension was assessed by administering two story-related questions after the recording session, with possible scores ranging from 0 to 2. Mean scores (± SD) were 1.9 ± 0.3 and 1.81 ± 0.4 for TD and DLD groups, respectively. A Wilcoxon rank-sum test revealed no significant difference between the groups (*z* = 0.45, *p* = 0.65), indicating that comprehension performance was comparable.

### 2.3. EEG data pre-processing

The EEG signals were first re-referenced to the mean of the mastoids. The EEG electrodes positioned at the jaw, mastoids, and forehead were excluded from the analysis, leaving 91 channels. The excluded channels were: E1, E14, E15, E17, E21, E32, E38, E43, E44, E48, E49, E56, E57, E63, E64, E68, E69, E73, E74, E81, E82, E88, E89, E94, E95, E99, E100, E107, E113, E114, E119, E120, E121, E125, E126, E127, and E128. Power line noise was removed using a notch filter. The data were then bandpass-filtered into specific frequency bands—delta (1-4 Hz), theta (4-8Hz), alpha (8-13 Hz), beta (13-25 Hz), low gamma (25-40 Hz) and whole frequency band (1-48 Hz) —using an 8^th^ order Butterworth filter. A zero-phase filtering method was applied to avoid phase shifts. The upper cutoff of 48 Hz was selected to remain below the 50 Hz power line frequency, thereby minimizing line noise contamination. Then the data was epoched to consecutive non-overlapping 5-second trials. A 5-second window was chosen to approximate time scales suitable for potential Brain Computer Interface (BCI) applications and to enable consistent segmentation across tasks for comparison. To remove noise sources like blinks, EOG or EMG signals in each trial, the channels with voltages over the absolute value of 100 μV were considered noisy and interpolated using the spline method. Trials were excluded if more than one-third of the total channels were deemed noisy. This automated threshold based procedure was selected to maintain a computationally efficient and fully automated pipeline compatible with potential real-time BCI applications, where manual inspection and ICA based cleaning are not feasible, and follows automated preprocessing approaches that use amplitude based thresholds for artifacts detection (Nolan et al., 2010; Jas et al., 2017). The data were then downsampled to 100 Hz to reduce computational load. After pre-processing, the average percentage of data removal or interpolated across all participants and frequency bands was 4.75%. On average, each participant retained 116.03 epochs (std=7.61) for the task.

### 2.4. Deriving unsupervised spatial filters using principal components analysis

A data-driven method was utilized to enable dimensionality reduction of the 91 channels and identify relevant spatial ensembles of channels that represented uncorrelated sources of cortical activity. In EEG data, signals recorded from different electrodes often share a large portion of variance due to volume conduction effects, where the same underlying neural sources are detected by multiple channels. This makes it challenging to isolate meaningful sources of brain activity by analyzing individual electrodes. Furthermore, EEG recordings represent indirect, large-scale measurements of neural activity, which are subject to estimating the original brain sources from observed signals. To address these challenges, we applied PCA to derive spatial filters that reduce dimensionality while preserving the dominant patterns in the signal.

To prepare the data, EEG epochs were preprocessed, standardized, and concatenated across trials. PCA was performed using singular value decomposition (SVD), which produced a set of principal components ordered by the amount of variance they explained. The contribution of each electrode to a given principal component was quantified by squaring the corresponding weights, since all squared weights for a component sum to one. This allowed us to interpret how strongly each channel contributed to each spatial pattern extracted by PCA. Three components explained almost 80% of the variance, and were retained for further analyses.

### 2.5. Principal component band power

To examine group differences in oscillatory dynamics during natural speech listening, theta/delta, low gamma/delta, and low gamma/theta band -power ratios were first computed from the PCA-derived spatial components. This was done because in the study reported by Araújo et al. (2024), the theta/delta power ratio was atypically high in the child *dyslexic* brain. This discovery led to the development of a successful BCI that reduced the theta-delta ratio during natural speech listening, leading to improvements in syllable stress processing and nonword decoding (Zheng et al., 2026). It was thus anticipated that excitation/inhibition ratios might also be important in distinguishing children with DLD with TD controls. After preprocessing, whole-brain EEG (91 channels) from all 5-s epochs was concatenated and subjected to PCA, retaining the first three components which together explained over 70% of the variance. Each epoch was projected onto these three spatial filters, and the power spectral density was estimated using Welch’s method with a single Hanning window (500 samples at 100 Hz). Delta- (1–4 Hz), theta- (4–8 Hz) and low Gamma- (25-40 Hz) power were extracted from the resulting spectra, and the band power ratio was computed for each epoch and each principal component. For each participant, the mean and variance of these per-epoch ratios across the entire story-listening session were used as summary metrics, providing indices of both the magnitude and the stability of power relationships in each spatial component.

### 2.6. Phase-amplitude coupling

Phase-amplitude coupling (PAC) measures whether low-frequency phases act as a temporal organizer for the amplitude of faster oscillations, thereby enabling different brain regions and networks to coordinate their activity across frequencies and locations. This neural mechanism is known to be important in adult cognition (e.g., learning and memory, Fries, 2015). PAC values hence represent hierarchical and instantaneous neural coordination across temporal scales. This cross-frequency interaction can be estimated from neural time series data by extracting the phase of the low-frequency signal and the amplitude envelope of the high - frequency signal, and measuring how strongly amplitude varies as a function of phase.

In the present study, PAC was evaluated using the Modulation Index (MI), a widely-used information-theoretic measure introduced by Tort et al. (2009). MI quantifies the deviation of the empirical phase–amplitude distribution from a uniform distribution. To compute MI, the phase of the low-frequency oscillation is divided into N phase bins, the mean high-frequency amplitude is calculated within each bin, and a discrete probability distribution 𝑃 over the phase bins is obtained by normalizing these amplitude values. The MI is then expressed as the Kullback–Leibler (KL) divergence between 𝑃 and the uniform distribution 𝑈:

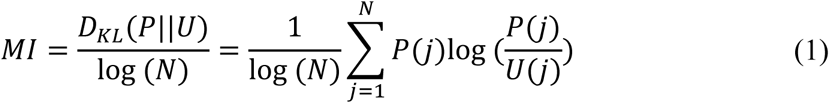

MI equals zero when amplitude does not depend on phase and increases towards one as phase–amplitude coupling becomes stronger.

PAC was computed for coupling between delta–theta (1–4 Hz phase; 4–8 Hz amplitude), delta–low-gamma (1–4 Hz phase; 25–40 Hz amplitude), and theta–low-gamma (4–8 Hz phase; 25–40 Hz amplitude) frequency ranges. PAC estimation was performed separately for the first three PCs (PC1–PC3) derived from each participant’s EEG response to continuous speech.

For each trial and PC, a comodulogram was generated to quantify PAC strength across the full phase and amplitude frequency grids, and the maximum z -scored MI (zMI) was extracted as a summary metric of coupling strength. For each participant, the mean and variance of trial -wise zMI values were then calculated for each PC, providing two participant-level summary measures (mean_zMI, var_zMI) used for group comparisons.

### 2.7. Deriving supervised spatial filters using common spatial pattern

While PCA is a powerful unsupervised method for identifying dominant spatial patterns in EEG data, it does not account for differences between experimental conditions or participant groups. To derive spatial filters that are specifically sensitive to group differences, we employed the CSP algorithm (Koles, 1991; Ramoser et al., 2000, Lotte et al., 2018, Duan et al., 2020). CSP is a supervised matrix decomposition technique widely used in EEG research to extract features that discriminate between two conditions or groups. Rather than maximizing variance generally (as PCA does), CSP identifies spatial filters that maximize variance in one class while minimizing it in the other class. This makes CSP especially useful for binary classification problems where the goal is to detect discriminative neural patterns.

More concretely, given a set of 𝑡 epoch segments 𝑋_𝑡_ ∈ ℝ^𝑑×𝑁^(where 𝑑 is the number of channels and 𝑁 is the number of datapoints on each epoch), epoch covariances 𝑋_𝑡_𝑋_𝑡_*^T^* ∈ ℝ^𝑑×^*^d^*, and Σ_1_ and Σ_2_ as the average epoch covariances for group 1 and group 2 subjects, CSP is calculated by the simultaneous diagonalization of the two average covariance matrices.

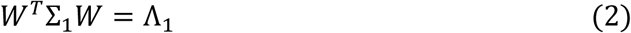

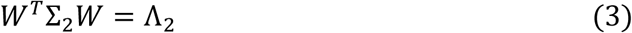

Where W is commonly determined so that Λ_1_ + Λ_2_ = 𝐼 (with Λ being a diagonal matrix of eigenvalues). Technically, this is achieved by solving the following generalized eigenvalue problem.

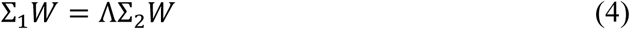

The spatially filtered signal S of this set of EEG epoch segments is then given by

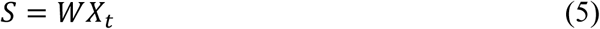

Assuming the column vectors in W are sorted by eigenvalue, its leftmost spatial filters (first column vectors) are maximizing the signal variance for group 1 and minimizing the signal variance for group 2, while the rightmost spatial filters (last column vectors) are maximizing the signal variance for group 2 and minimizing the signal variance for group 1.

### 2.8. Linear classifier

To perform supervised classification based on the spatial features extracted by the CSP algorithm, we used a Support Vector Machine (SVM). After projecting each EEG trial 𝑆 = 𝑊𝑋_𝑡_ onto the selected CSP spatial filters 𝑊, we computed the log-variance of the resulting time series to generate features for classification.

Specifically, for each CSP component 𝑆_𝑖_, the log-variance was calculated as:

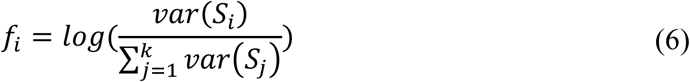

Where k is the total number of selected components. This feature transformation ensures comparability across trials and improves the distributional properties of the data.

A linear SVM was then applied to these features to learn a decision boundary between the two groups. The SVM algorithm identifies the hyperplane that best separates the classes with the largest possible margin in the feature space. As a supervised learning method, SVM is well-suited for EEG classification and is commonly used in brain-computer interface research. In this context, SVM served as the final step in supervised spatial filtering pipeline, following CSP feature extraction, to enable discriminative classification of neural patterns.

### 2.9 Statistical Approach

This was modelled closely on Araújo et al. (2024), who applied a similar pipeline to story listening data collected from children with either developmental dyslexia (N = 16) or DLD (N = 7) and TD control children (N = 40).

#### 2.9.1 Principal Components Analyses (PCA)

First, in order to explore the general neural patterns associated with story listening and to understand how different frequency bands contribute to these processes, we applied PCA across frequency bands (delta, theta, alpha, beta, low gamma). Our goal was to identify key spatial patterns of neural activity and determine whether distinct frequency bands reveal unique neural dynamics during story comprehension. We conducted both separate group comparisons and a combined analysis of EEG data from both participant groups, and assessed the variance explained by principal components (PCs) to uncover any group-specific differences.

#### 2.9.2 Band Power Ratios and Phase-Amplitude Coupling analyses

Second, to characterize potential group differences in oscillatory power relationships and cross frequency coupling, statistical analyses were performed to assess group differences in both band power ratios and PAC metrics. For each outcome measure, normality of the participant-level distributions was assessed using the Shapiro–Wilk (SW) test. Depending on the outcome of these tests, comparisons between the DLD and TD groups were carried out using either independent-samples t-tests (for normally distributed data), or Mann–Whitney U (MWU) tests (for non-normal data).

For band-power ratios, statistical comparisons were conducted separately for each of the three principal components (PC1–PC3) and for each frequency-band pairing (theta/delta, theta/low-gamma, delta/low-gamma). Participant-level metrics included the mean and variance of epoch-wise ratios.

For PAC, group comparisons were performed on participant-level mean zMI and variance zMI for each PC and coupling type (delta–theta, delta–low-gamma, theta–low-gamma). Resulting *p*-values were corrected for multiple comparisons using the false discovery rate (FDR) procedure.

In addition to statistical comparisons of summary metrics, polar plots were generated to visualize the spatial and phase-specific characteristics of PAC at the group level.

#### 2.9.3 Common Spatial Pattern and Classifier analyses

Third, in order to determine whether distinct oscillatory dynamics during natural speech listening could be robust enough to build a classifier for identifying DLD, we applied CSP to identify biomarkers that might differentiate the groups across all frequency bands. Each CSP calculation produced the same number of filters as the number of recorded channels. Following common practice in BCI research, only a subset of these filters was analyzed. For each group comparison, the two filters that maximized variance for each group (m = 2; total number of filters = 4) were computed. To explore potential group differences in spatial patterns within each frequency band, group differences were then assessed. Finally, to test the robustness of the CSP-based features for classification, we applied a linear SVM approach to differentiate between the two participant groups across all frequency bands. We used a cross-validation approach, in which data from two participants from each group (out of 16 participants per group) were left out to form the test set, resulting in four participants being held out at a time. The data from the remaining 28 participants were used as the training set. This process was repeated randomly for 90 iterations, rotating the combination of test participants from a total of 14400 possible combinations 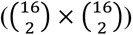. By using repeated assessments of the model’s performance, we minimized the influence of individual differences between participants and maximized the use of available data for training, leading to more robust and reliable classification results. Within the training set, we performed 4-fold cross-validation to optimize the hyperparameters. After determining the best hyperparameter configuration, we trained the final SVM model using all 28 participants in the training set and evaluated its performance on the test set. We assessed the model’s performance in three ways: First, by comparing the overall accuracy on the test set. Second, by calculating the accuracy for each participant as the ratio of correctly classified trials, then determining how many participants achieved an accuracy greater than 50%. Third, we computed the Area Under the Curve (AUC) based on the accuracy of each participant. In addition, we computed correlation maps in order to visualize the key neural regions contributing to the group differences.

#### 2.9.4 Correlation-Based Scalp Mapping of Spatial Features

Fourth, we identified the spatial features that most strongly differentiated the DLD group from the age-matched control group. These features may reflect brain regions showing atypical activation in DLD participants compared to controls. To achieve this, we correlated the filtered data from each filter with the original EEG data from each channel. By projecting these correlations onto the scalp, we were able to assess how much each channel contributed to the filtered data, essentially determining the strength of the information encoded by each channel in the filtered signals.

#### 2.9.5. Integration of PCA and CSP Spatial Patterns

Finally, to assess the stability and relevance of spatial features identified by our analyses, we compared the topographic projections derived from PCA and CSP. PCA, as an unsupervised method, identifies components that capture the greatest variance in the data without considering group membership, whereas CSP, as a supervised method, emphasizes variance differences between groups by extracting spatial filters that maximize variance in one class while minimizing it in the other. The detection of similar spatial patterns in both PCA and CSP suggests that features carrying high variance in an unsupervised manner may also contribute critically to distinguishing the two groups. Overlap between PCA and CSP spatial patterns therefore indicates that the neural sources explaining the greatest variance during story listening are also those most relevant for differentiating children with DLD from their typically developing peers. Such convergence strengthens confidence that the identified patterns are not artifacts of a single method, but rather reflect stable neural processes with both cognitive significance and clinical significance.

To quantify the shared spatial information, we first calculated channel-wise correlations between each PC time series and the original EEG channels, producing a scalp projection for each component. The same procedure was applied to each CSP-filtered signal, resulting in corresponding CSP scalp projections (CSP1, CSP2, CSP90, CSP91). Cosine similarity was then calculated between the correlation maps of the two methods within each group and frequency band. For the Control group, similarities were computed between the three PC correlation maps and CSP1 and CSP2, whereas for the DLD group, similarities were computed between the three PC correlation maps and CSP90 and CSP91. These correlation maps (one value per electrode) were then compared pairwise between the three PCs and the two CSP filters within each frequency band and group. The cosine similarity was used to quantify the global similarity between each pair of maps. The overlap maps were generated using the absolute method, where the absolute values of the PCA and CSP correlation maps were multiplied pointwise to capture regions of strong shared contribution irrespective of polarity.

## 3. Results

### 3.1. PCA Analysis Across Frequency Bands

EEG data from both groups of participants were first combined and subjected to PCA in each frequency band. PCs with the highest eigenvalues were retained, setting the threshold at a total of 70% variance explained. Based on this criterion, the first three PCs were retained in all frequency bands. These preliminary analyses are available as Supplementary Section 1.

### 3.2. PCA Analysis by Participant Group

Of greater theoretical interest regarding TS theory, we next ran the same PCA analyses by group. These analyses revealed discrepancies in both the delta and theta bands for both PC2 and PC3 for the children with DLD, as shown in Figure 1. Discrepancies in the other bands did not occur, please see Supplementary Figure S2. Inspection of Figure 1 reveals that while the TD control children show highly similar PCA topography and weightings to the larger group of TD children analysed by Araújo et al. (2024, see Figure 1B), the children with DLD in the current study appear to show a reversal of PC2 and PC3 in the delta band compared to the current TD control group. Visually, this reversal appears to be ameliorating in the theta band, although some extra right lateralized weights are visible. Inspection of Figure 1 further suggests that PC2 in the control children in the delta band corresponds to PC3 for the children with DLD, while PC3 in the control children in the delta band corresponds to PC2 for the children with DLD. While PC2 and PC3 together account for ∼41% of variance for the TD group in the delta band regarding neural activation during story listening, for the DLD group they account for only ∼30% of variance. For the DLD group, more variance is accounted for in the delta band by PC1 (∼50%) compared to TD controls (∼40%), and this is still the case in the whole band analyses (1 – 48 Hz; DLD PC1 ∼49%, TD PC1 ∼39%). This could suggest less efficient utilization of similar speech processing mechanisms by the DLD brain in the delta and theta bands. In order to inspect this possibility further, power ratio and phase-amplitude coupling analyses were run by group.

**Figure 1.**
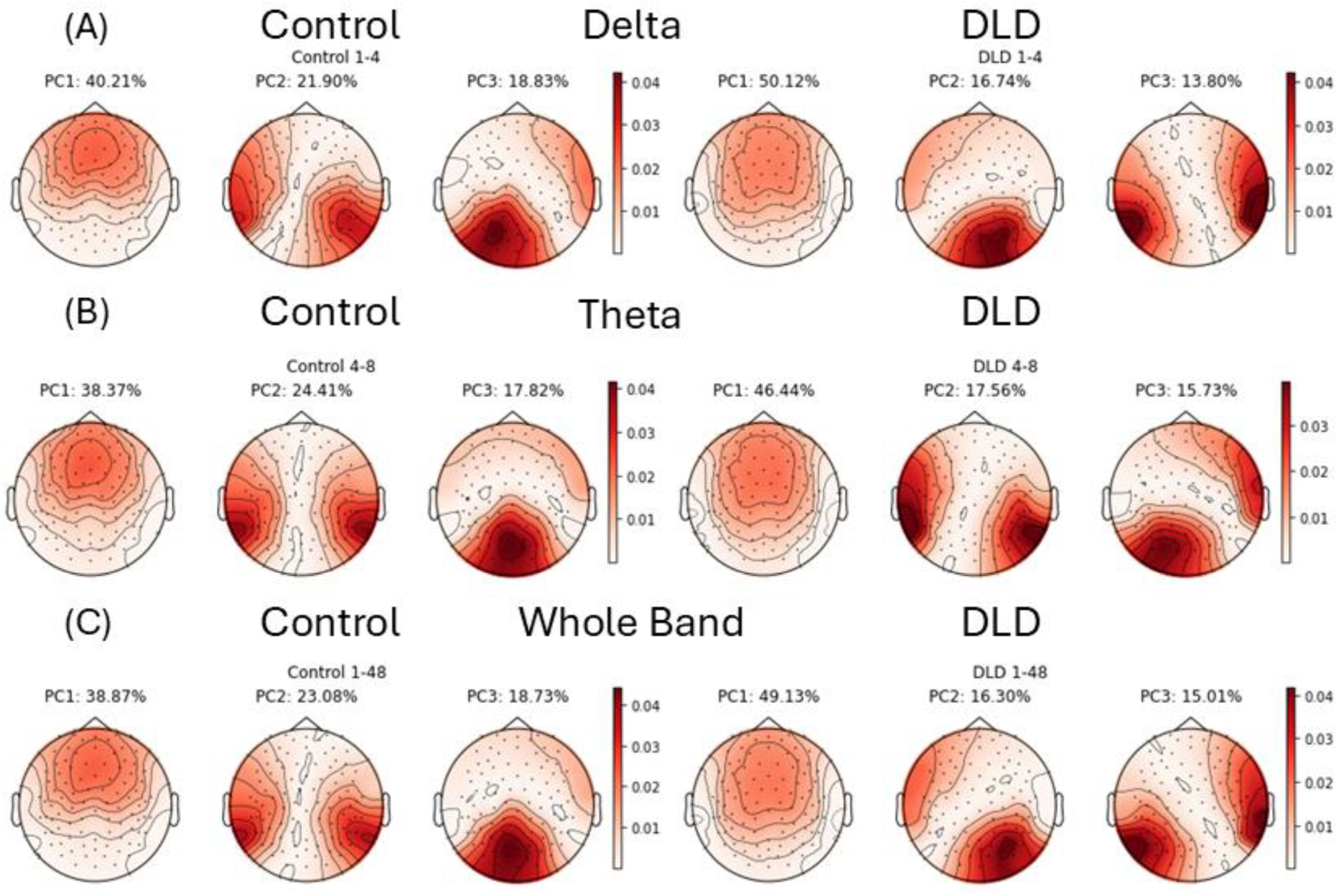
Comparison of PCA Results by Group for the Delta and Theta bands and Whole Band. (A) Delta band, (B) Theta band, and (C) whole frequency band results. For each band, the first three principal components (PC1–PC3) are shown separately for the Control group (left) and the DLD group (right). The percentage above each map indicates the variance explained by that component. Topographic maps represent the spatial distribution of component loadings across electrodes.

### 3.3. Principal Component Band Power and PAC Analyses

First, the power ratio between pairs of frequency bands (theta/delta, theta/low gamma, and delta/low gamma) was computed to assess whether there were any differences in power ratios between groups (see Section 2.5 and 2.9.2). Power spectral estimates were obtained from the EEG data projected onto the first three principal components (PCs) derived from each participant’s PCA decomposition. Across all frequency pairs and PCs, no significant group differences were identified. The full set of power-ratio comparisons is reported in Supplementary Table S1. Accordingly, in contrast to the child dyslexic brain, there are no apparent excitation/inhibition imbalances during natural speech listening in the child DLD brain. However, there may still be differences in how low-frequency phases affect oscillatory power.

To explore potential group differences in cross-frequency coupling during continuous speech listening, we examined two complementary PAC outcome measures: (1) the distribution of participant-level PAC metrics (mean and variance of trial-wise zMI, see section 2.9.2), and (2) polar plots summarising the preferred phase of coupling. Polar plots illustrate the phase angle of the low-frequency oscillation at which high-frequency amplitude is maximal, allowing comparison of preferred phase patterns between groups. Together, these measures provide a comprehensive depiction of potential PAC differences between TD children and those with DLD across the first three spatial components of the EEG response.

Figure 2 presents the participant-level distributions and polar plots for the components which showed significant group differences before FDR correction. These group differences occurred in PC2 in delta–theta PAC and PCs 1 and 2 in delta–low gamma PAC. Shapiro-Wilk (SW) analyses showed that data in each set of analyses met the conditions for normality, with a few exceptions for which the group comparisons utilised the non-parametric Mann-Whitney U test. The SW analyses are reported in Supplementary Table S2.

**Figure 2.**
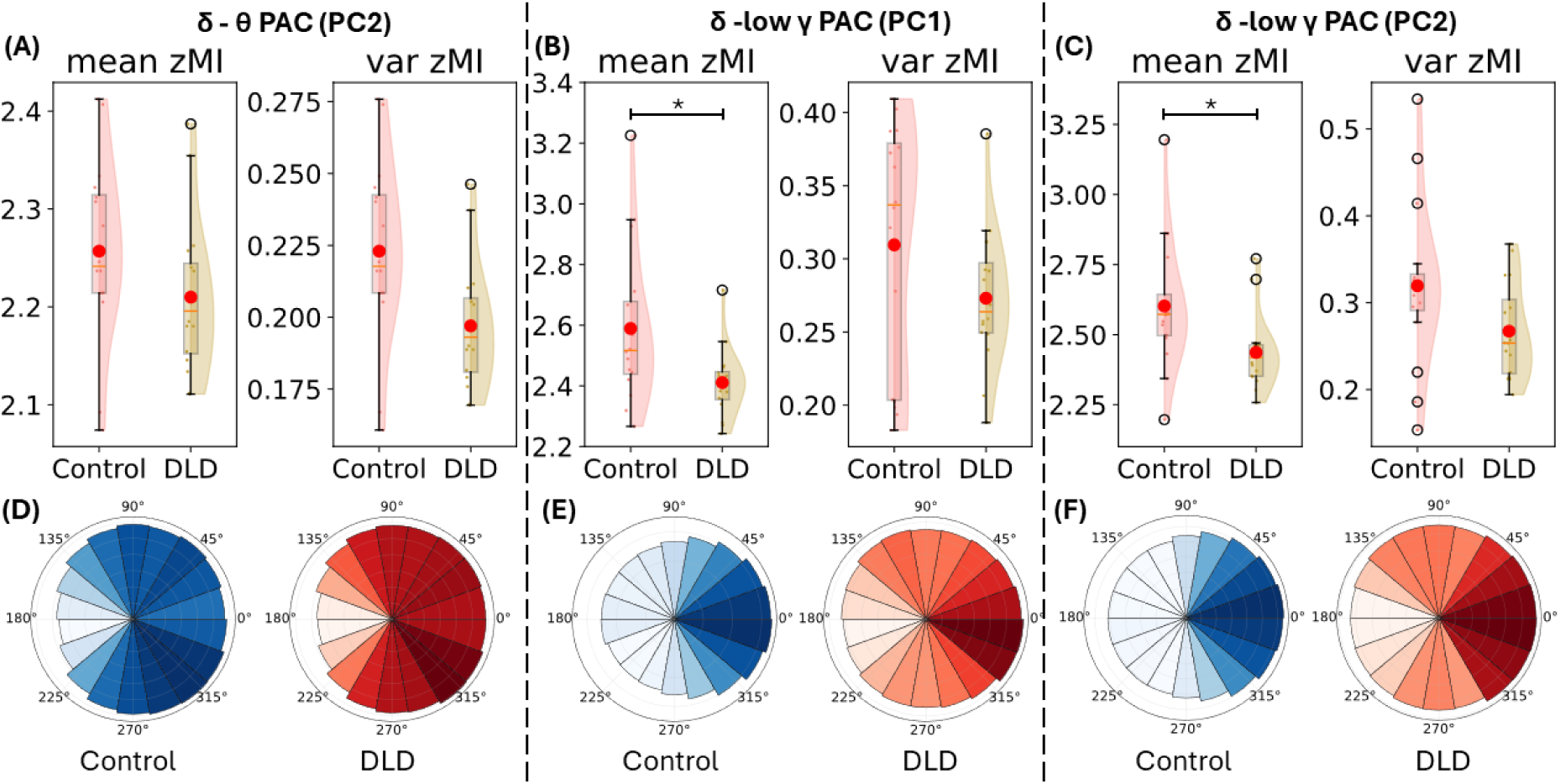
Participant-level distributions for delta-driven PAC and polar plots for mean PAC in each case. Panels (A) and (D) correspond to delta–theta PAC for PC2, Panels (B) and (E) correspond to delta–low gamma PAC for PC1, and Panels (C) and (F) correspond to delta–low gamma PAC for PC2.

Considering each analysis in turn:

1) Delta–theta PAC

For delta–theta PAC, a group difference was detected in the variance of zMI on PC2. The children with DLD showed lower variability of coupling strength compared to TD controls (uncorrected *p*=0.013]). However, this effect did not survive FDR correction, and no other significant delta–theta differences were observed for PC1 or PC3. Accordingly, the atypical low-frequency PAC results reported by Araujo et al. (2024) were not supported. Delta–theta PAC metrics are shown in Table 2.

**Table 2.** Participant-level Delta-Theta PAC metrics for TD and DLD groups (mean and variance of zMI; corrected *p* values)

| Data | group | mean | STD | p-value<br>(MWU test / t-test) |
| --- | --- | --- | --- | --- |
| PC1 - mean | Control | 2.280 | 0.063 | 0.255(t-test) |
|  | DLD | 2.323 | 0.101 |  |
| PC1 - var | Control | 0.220 | 0.023 | 0.216(t-test) |
|  | DLD | 0.249 | 0.056 |  |
| PC2 - mean | Control | 2.257 | 0.094 | 0.255(t-test) |
|  | DLD | 2.210 | 0.079 |  |
| PC2 - var | Control | 0.223 | 0.032 | 0.078(t-test) |
|  | DLD | 0.197 | 0.022 |  |
| PC3 - mean | Control | 2.248 | 0.092 | 0.925(MWU test) |
|  | DLD | 2.238 | 0.060 |  |
| PC3 - var | Control | 0.206 | 0.028 | 0.491(t-test) |
|  | DLD | 0.216 | 0.038 |  |

2) Delta–low gamma PAC

By contrast, delta–low gamma PAC revealed statistically robust group differences following FDR correction. Significant effects were detected in the mean zMI values for PC1 and PC2, with children with DLD showing reduced delta–low gamma coupling compared to TD controls (corrected *p*: 0.032 and PC1 and 0.024 for PC2). As the data failed the normality assumption for children with DLD in PC2, the Mann-Whitney U test was used. The data indicate reliable group differences in delta-low gamma PAC that generalised across both central (PC1) and temporal (PC2) spatial components. Trial-wise zMI variance did not differ significantly between groups for any PC. Delta-Low Gamma PAC metrics are shown in Table 3.

**Table 3.** Participant-level Delta-Low Gamma PAC metrics for TD and DLD groups (mean and variance of zMI, corrected *p* values)

| Data | group | mean | STD | p-value<br>(MWU test / t-test) |
| --- | --- | --- | --- | --- |
| PC1 - mean | Control | 2.589 | 0.248 | **0.032**(t-test) |
|  | DLD | 2.411 | 0.110 |  |
| PC1 - var | Control | 0.310 | 0.083 | 0.147(MWU test) |
|  | DLD | 0.273 | 0.046 |  |
| PC2 - mean | Control | 2.602 | 0.225 | **0.024**(MWU test) |
|  | DLD | 2.436 | 0.129 |  |
| PC2 - var | Control | 0.319 | 0.092 | 0.091(t-test) |
|  | DLD | 0.267 | 0.054 |  |
| PC3 - mean | Control | 2.543 | 0.211 | 0.617(MWU test) |
|  | DLD | 2.485 | 0.120 |  |
| PC3 - var | Control | 0.296 | 0.069 | 0.198(t-test) |
|  | DLD | 0.268 | 0.061 |  |

3) Theta–low gamma PAC

No significant group differences were detected for theta–low gamma PAC and accordingly the analyses are not reported.

### 3.4. CSP-based classification of DLD and TD control group using oscillatory dynamics

As a sanity check prior to running the CSPs, we calculated the power spectral density (PSD) of the filtered data and compared the statistical differences in variance between the two groups for each filter across all frequency bands. The results are presented in Figure 3 (A). The PSD analysis confirmed that the filters functioned as intended: they maximized the variance in the target group while minimizing it in the other group. Using the Mann-Whitney test, we found that the variance of the filtered EEG data between the two groups was significantly different across almost all frequency bands (Bonferroni corrected p-value < 0.05), with the exception of the second filter (CSP2) in the control group for the alpha band, and the first filter (CSP90) for the DLD group in the theta band, which did not reach significance. This suggests that the CSP-determined filters were largely successful in separating the two groups. A representative example is depicted in Figure 3 (B). While the TD group shows a left-lateralised spatial pattern, the DLD group shows an atypical right frontal spatial pattern. A full depiction of all CSPs is provided as Supplementary Figure S3. Inspection of Figure S3 shows atypical right frontal spatial patterns for the DLD group in every oscillatory band, although at higher frequencies one of the two DLD CSPs typically shows the TD left-lateralised spatial pattern.

**Figure 3.**
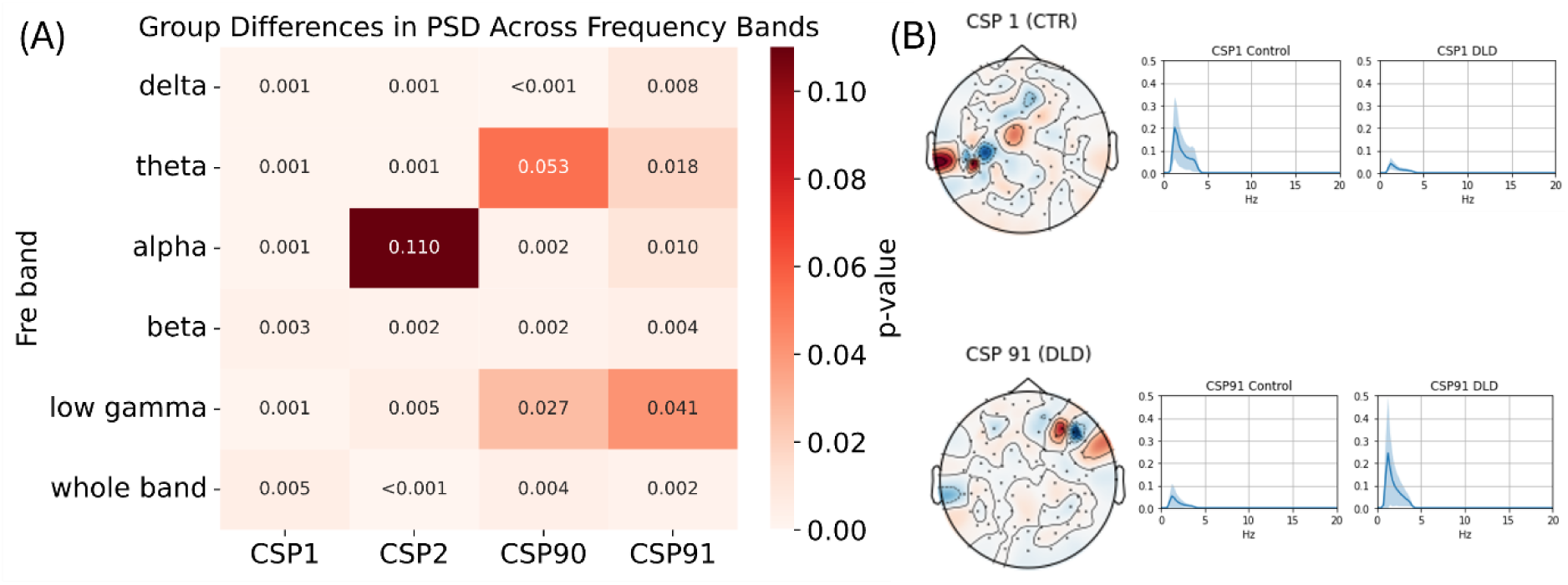
PSD Differences in CSPs Across Frequency Bands. Panel (A) shows the Bonferroni-corrected comparisons, Panel (B) shows an example of a filter separating the groups in the delta band.

### 3.5. Training a Linear Classifier with Features based on CSP filters

To test the robustness of CSP-based features for group classification, we applied a linear SVM as detailed in Sections 2.8 and 2.9.3. The model’s performance across different frequency bands is presented in Table 4 and Figure 4. As shown in Table 4, we observed an average test accuracy ranging from 0.665 to 0.744, a participant accuracy between 0.644 and 0.833, and AUC ranging from 0.715 to 0.855. Figure 4 shows that all CSPs enable above-chance classification. These results suggest that the spatial filters applied to the story-listening EEG data encode valuable information for identifying DLD participants compared to age-matched controls.

**Figure 4:**
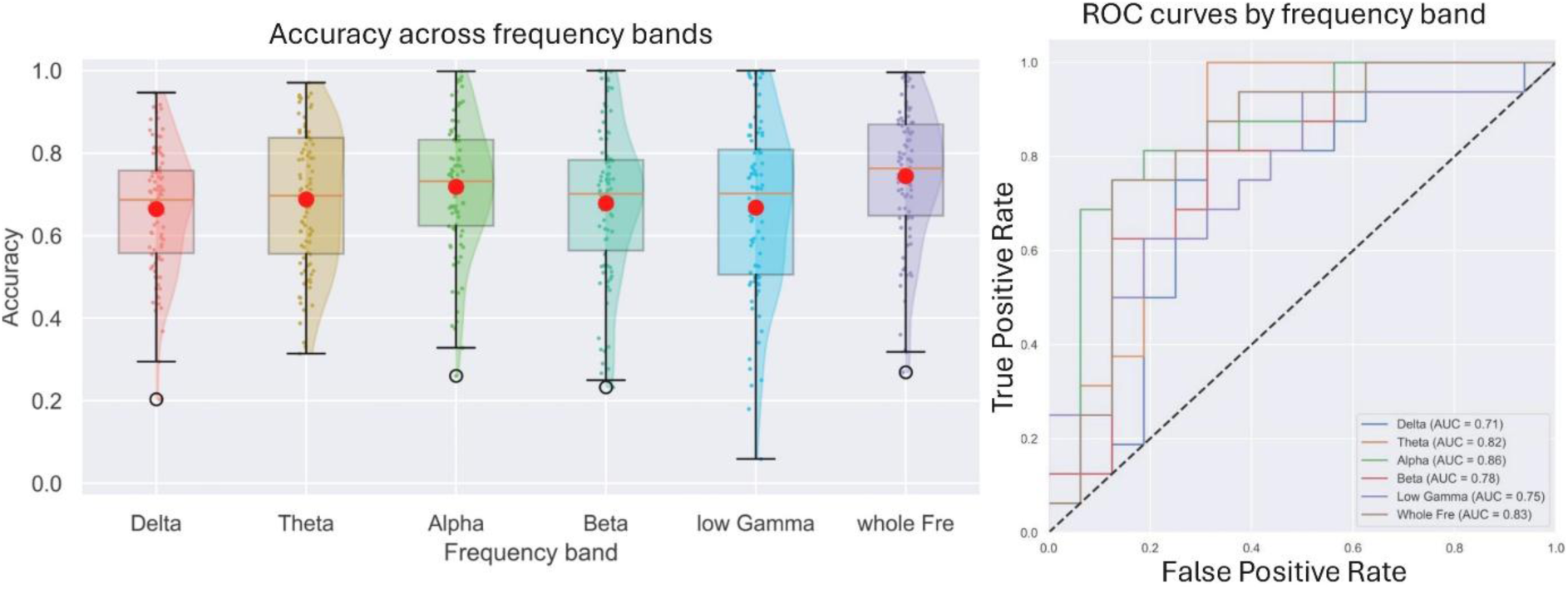
SVM classification performance across frequency bands. Left panel shows the distribution of classification accuracy across frequency bands, with boxplots indicating median and interquartile range and red dots marking mean accuracy. Right panel shows ROC curves for each frequency band, with AUC values listed in the legend.

**Table 4.** Performance of the SVM models by frequency band.

| Fre Band | best validation score | test score | participant accuracy | AUC |
| --- | --- | --- | --- | --- |
| Delta | 0.612 | 0.665 | 0.767 | 0.715 |
| Theta | 0.643 | 0.688 | 0.756 | 0.820 |
| Alpha | 0.646 | 0.719 | 0.811 | 0.855 |
| Beta | 0.680 | 0.679 | 0.706 | 0.777 |
| Low Gamma | 0.669 | 0.668 | 0.644 | 0.752 |
| Whole Fre | 0.687 | 0.744 | 0.833 | 0.828 |

### 3.6 Spatial Feature Identification of DLD and Control Group

After observing robust classification performance, we computed channel-CSP correlation maps to visualize the key neural regions contributing to the group differences. These maps reflect the correlation between the CSP-filtered signals and the original channel data, allowing us to identify brain regions showing atypical activation in DLD participants compared to TD controls. The resulting topographic correlation maps are presented in Figure 5. The filters CSP1 and CSP91 were selected for this purpose by the CSP algorithm because they maximized variance more strongly for the TD versus DLD groups compared to other CSP filters (clearly visible in Supplementary Figure 3). Supplementary Figure 3 displays the projections of the CSP filters themselves. The spatial patterns of CSP2 and CSP90 were less consistent across frequency bands. In the delta band, CSP1 (maximizing TD variance) showed pronounced contributions from the central, left temporal, and right parietal regions, whereas CSP91 (maximizing DLD variance) exhibited dominant weights over the frontal and central areas. This pattern contrasted with the other frequency bands (theta, alpha, beta, and low gamma), where CSP1 consistently displayed bilateral temporal and bilateral parietal distributions, and CSP91 showed stronger contributions from the left parietal and left central regions. Together, these findings suggest that group-specific spatial dynamics are most pronounced in the delta band, with control participants showing broader engagement of parietal and temporal regions and DLD participants exhibiting stronger central and frontal activity. Accordingly, the CSP analyses pinpoint the delta band as a key mechanistic source of group differences.

**Figure 5:**
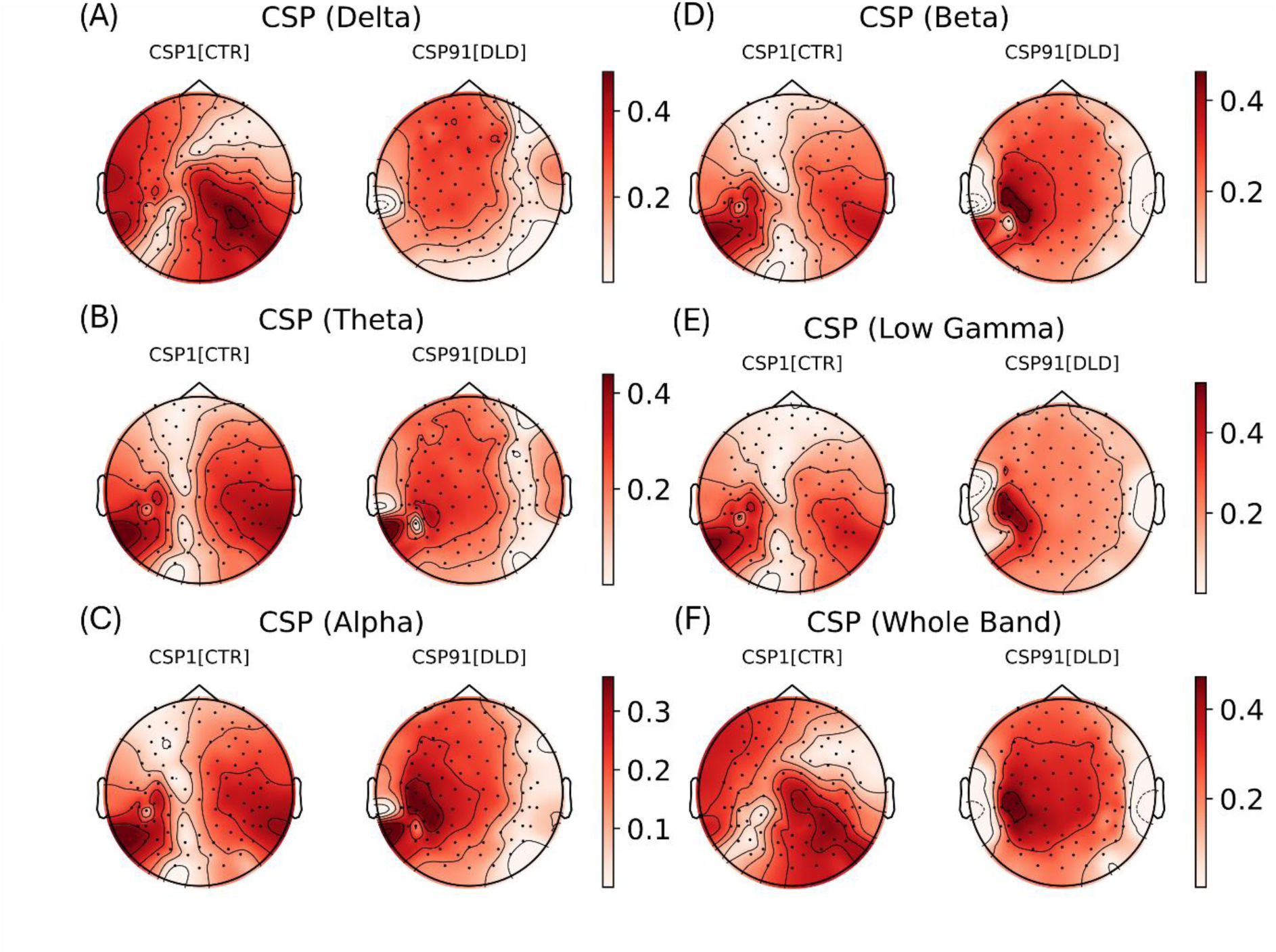
Channel–CSP correlation topographies for CSP1 and CSP91 across frequency bands and both groups. (A) Delta, (B) Theta, (C) Alpha, (D) Beta, (E) low Gamma, and (F) whole frequency band. For each band, correlation maps are shown for CSP1 (maximizing variance for the Control group; left) and CSP91 (maximizing variance for the DLD group; right). The topographies represent the correlation between the CSP-filtered signals and the original channel data, indicating the relative spatial contribution of each electrode to the discriminative components.

### 3.7. Correlational analysis of PCA and CSP spatial features in distinguishing group differences

It can be observed by comparing Figures 1 and 5 that when both the PCA channel weights and the CSP correlation weights are projected onto the scalp, some of the regions highlighted in the CSP correlation maps resemble the patterns seen in the PCA channel weight projections. While PCA maximizes variance locally by identifying components that capture the most variance, CSP emphasizes the variance differences between two groups, seeking features that maximize variance in one class while minimizing it in the other. The detection of potentially similar patterns in both PCA and CSP suggests that features carrying high variance in an unsupervised manner may also play a key role in distinguishing the two groups. If similar spatial patterns are observed across PCA and CSP, this would indicate that neural sources explaining the largest proportion of variance (as identified by PCA) also contribute to group discrimination (as identified by CSP). Such correspondence would suggest that the variance structure of the EEG data inherently encodes group-related differences. Further, the convergence of spatial patterns across the two methods would strengthen the interpretability of the findings, indicating that these regions likely represent stable neural sources relevant both to general neural processing during story listening and to group-specific characteristics that distinguish DLD from TD participants.

As a statistical test of these observations, we examined the spatial correspondence between PCA- and CSP-derived features by correlating each EEG channel with the top three principal components and with the CSP-filtered signals (CSP1, CSP2, CSP90, CSP91; see Section 2.9.5), and quantified the degree of spatial similarity using cosine similarity. For the TD control group, the comparison involved the three PC correlation maps and CSP1/CSP2, whereas for the DLD group, comparisons were made with CSP90/CSP91. The overlaps with absolute cosine similarity greater than 0.7 differed by group, and are illustrated in Figure 6. All overlap maps are provided in Supplementary Figure S4.

**Figure 6:**
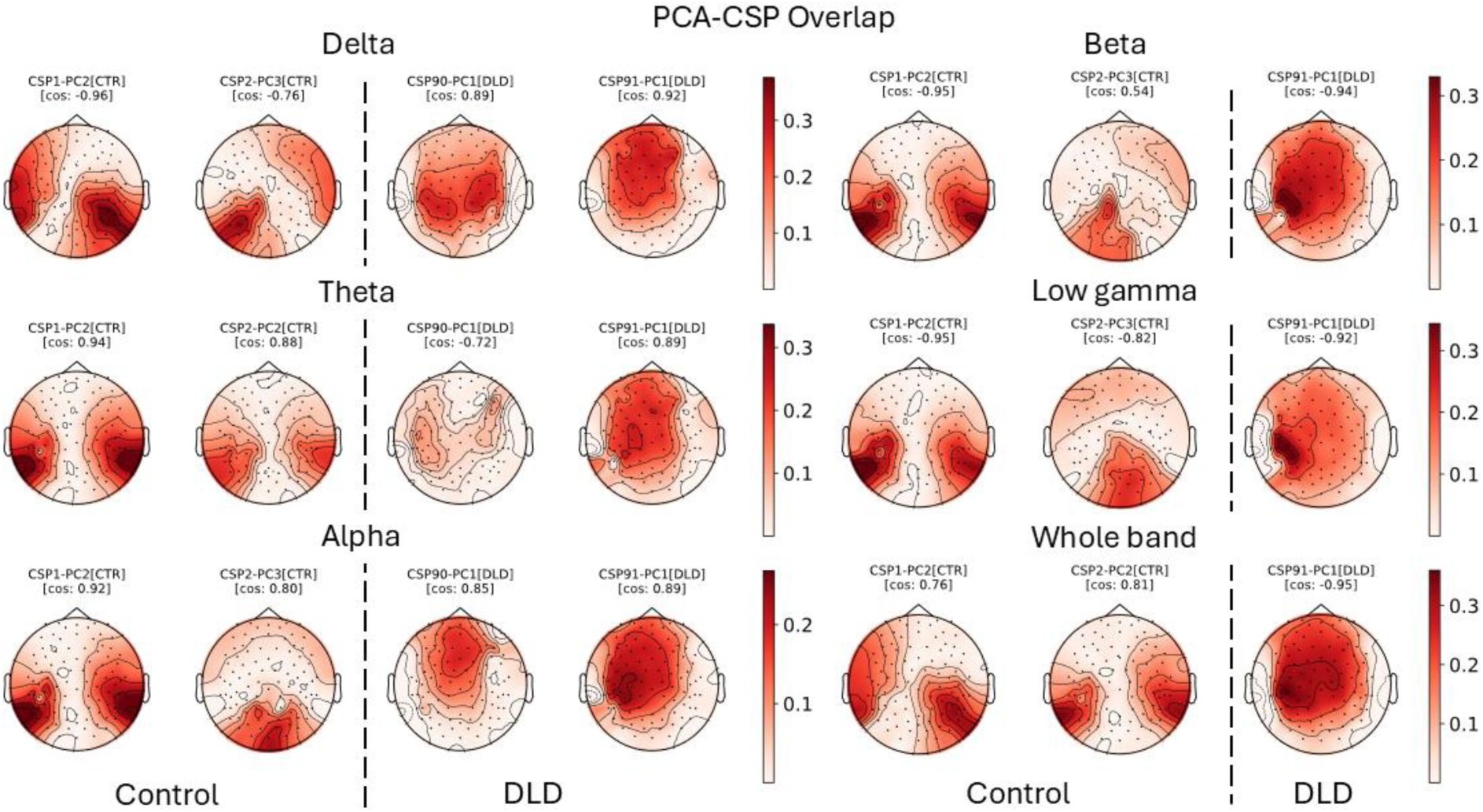
Spatial overlap between PCA and CSP across frequency bands . Topographic maps show the spatial correspondence between PCA and CSP derived correlation patterns across frequency bands (delta, theta, alpha, beta low gamma, and whole band). For the TD group (left of dashed line), overlaps are shown between the top three PC correlation maps and CSP1/CSP2. For the DLD group (right of dashed line), overlaps are shown between PC maps and CSP90/CSP91.

Inspection of Figure 6 shows that PC1 was the primary driver of consistency for the DLD brain (rightward panels for each frequency band), whereas PC2 and PC3 were the drivers of consistency for the TD brain (leftward panels for each frequency band). Distinct spatial organization was thus observed between groups. In the TD control group, overlap regions were mainly located in the bilateral temporal and parietal areas across the delta, theta, and beta bands, and shifted toward the bilateral parietal and occipital regions in the alpha and low gamma bands. A similar bilateral temporal–parietal pattern was observed when considering all bands together (1-48 Hz). In contrast, the DLD group exhibited overlap primarily in the central and frontal regions across the delta, theta, and alpha bands, with left lateralized temporal activation appearing in the alpha band and getting stronger in the faster beta and low gamma bands, which exhibited mainly central activity. In the whole-band analysis, the DLD group again showed overlap concentrated in the central and frontal regions, with some left lateralization. Accordingly, the spatial overlap for the two analysis methods was greater for PC2 and PC3 for the TD children, and for PC1 for the DLD children.

## Discussion

The PCA, PAC and CSP analyses presented here indicate that atypical automatic low-frequency neural oscillatory responses to natural speech can uniquely identify children with DLD. The PCA analyses identified the delta and theta oscillatory bands as the source of group differences, and this focus on low-frequency neural responses was supported by the PAC analyses. Contrary to prediction, however, it was delta-low gamma PAC that was atypical in the children with DLD, not delta-theta PAC as identified in the study by Araújo et al. (2024). Interestingly, theta-gamma PAC, which is known to be a significant predictor of language acquisition by infants (Attaheri et al., 2024) and is also critical for speech processing by adults (López-Madrona et al., 2025), did not differ by group. As Araújo et al. (2024) only included 7 participants with DLD, the current findings regarding delta-low gamma PAC should be considered more reliable. In the infant studies reported by Attaheri et al. (2022, 2024), both delta-low gamma PAC and theta-low gamma PAC to continuous speech were significant at each age measured (4, 7 and 11 months). In adult studies, delta-low gamma PAC is rarely measured (Fontolan et al., 2014), so its potential role in expert speech processing is unclear. Nevertheless, the PAC data from infants and children suggest that *both* theta-low gamma and delta-low gamma PAC mechanisms may contribute to the initial foundation of a language system.

The current data thus support the claim that low-frequency (i.e., delta and theta) oscillatory mechanisms are fundamental to understanding the aetiology of DLD, as predicted by TS theory (Goswami, 2011, 2022). The CSP analyses presented here provided further support for this low-frequency oscillatory perspective. Although the spatial filters that maximised the variance for one group and minimised it for the other group were significantly different in virtually all frequency bands examined (see Figure 3), the classifier analyses based on the CSP filters identified the delta band as the key source of group differences (Figure 5). When the convergence of the spatial patterns identified by the PCA and CSP analyses was computed, the main overlap for the TD group was in bilateral and temporal areas across the delta and theta bands, but the main overlap for the DLD group was in the central and frontal areas for the delta and theta bands. These data suggest that the topography of the processing of low-frequency speech information differs in TD and DLD children and can be a useful basis for classification.

Regarding the wider DLD neural literature, it is difficult to draw direct comparisons with the data reported here as very few prior studies have adopted an oscillatory perspective on continuous speech encoding. A recent MEG study using single words as speech input reported that Finnish children with DLD showed poorer cortical representation of the amplitude envelope information compared with TD children at longer latencies (at ∼200–300 ms lag) (Nora et al., 2024). This appears consistent with the TS-oscillatory perspective adopted here. A recent TS-driven EEG study using rhythmic speech input (audio-visual presentation of the syllable ‘ba’ repeated at 2Hz) also found atypical cross-frequency dynamics for the children with DLD. For rhythmic speech, phase-phase coupling between low-frequency phases (both delta and theta) and low-gamma phases appeared to operate differently in the children with DLD compared to control children (Keshavarzi et al., 2024). As also noted previously, studies of toddlers and young children (16-, 24- and 36-month-olds) who are at family risk for DLD have reported that resting state power in the gamma band is associated with later language outcomes (Gou et al., 2011), while infant at-risk studies measuring resting state theta power have not found significant prediction of later language outcomes (Cantiani et al., 2016). Nevertheless, differences in resting state power may not be indicative of how the speech processing system performs when exposed to continuous speech. Most neural studies of children with DLD listening to *continuous* speech have utilized fMRI, and these studies typically show reduced activation in left frontal and temporal cortical areas during language processing in the DLD group (Asaridou and Watkins, 2022, for review). Accordingly, these fMRI data are consistent with the spatial data presented here.

In conclusion, here we identify a range of low-frequency neural processing patterns that are atypical in children with DLD. These data support the utility of employing a TS framework to explain developmental disorders of language learning. The findings are suggestive of distinct atypical neurocognitive speech encoding mechanisms underlying DLD focused on extracting prosodic information and syllabic rhythm patterns, which could be targeted by novel interventions.

## ACKNOWLEDGEMENTS

The authors would like to thank all the participants who volunteered for the study. This research was funded by a donation to U.G. from the Yidan Prize Foundation. The sponsor played no role in the study design, data interpretation, nor writing of the report.

## DATA AND CODE AVAILABILITY

Data and code will be made available on request.

## DECLARATION OF COMPETING INTEREST

The authors declare no conflicts of interest.

XZ – Conceptualisation, Data Curation, Methodology,

Data Analysis, Visualisation, Writing – original draft

JA – Conceptualisation, Supervision, Methodology,

Writing – Review and editing

MK – Investigation, Data Curation,

Writing – Review and editing

GF – Investigation,

Writing – Review and editing

SR – Investigation,

Writing – Review and editing

LP – Investigation,

Writing – Review and editing

UG - Conceptualisation, Project Administration, Funding Acquisition, Resources, Supervision,

Writing – original draft

