## Supplementary materials for "Atypical cortical encoding of the low-frequency temporal dynamics of natural speech identifies children with Developmental Language Disorder"

*1. PCA Analysis Across Frequency Bands: All Children Combined*

The channel weights of the first three PCs were projected onto the scalp to investigate their spatial distribution, which is shown in Figure S1. The variance explained by each PC is also indicated. PC1 (total variance explained across all bands: 39.56%-44.96%) showed strong channel weights in the central and frontal regions of the scalp. Spatially this component was very similar to the first PC identified in the natural speech listening data analysed by Araújo et al. (2024, Figure 1), which explained 49.9% of the variance in that study. PC2 (total variance explained across all bands: 17.33%-22.23%) typically showed channel weights in the bilateral temporal regions, with the exception of the delta band. In our previous dataset (Araujo et al., 2024), PC2 also showed weights in bilateral temporal regions, and accounted for 14.5% of the variance. In the current dataset, the increased variance in the delta band appeared to be driven by group differences, described in the main paper. PC3 (total variance explained across all bands: 13.27%-17.30%) mostly showed channel weights in the parietal and occipital regions, again matching the PC3 identified in our previous dataset (Araújo et al., 2024), where it accounted for 12.8% of the variance.

**Figure S1.** PCA results combining across group by band

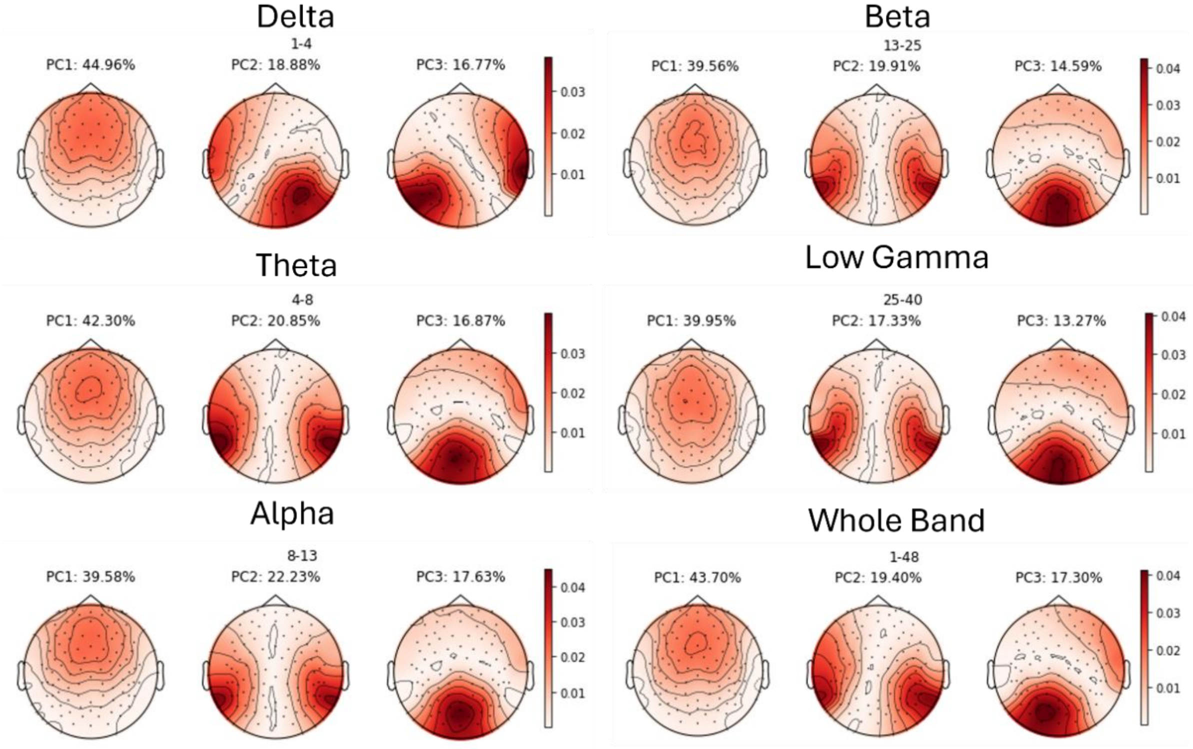

2. ***Figure S2.*** *Group-Specific PCA Analyses Across Frequency Bands*

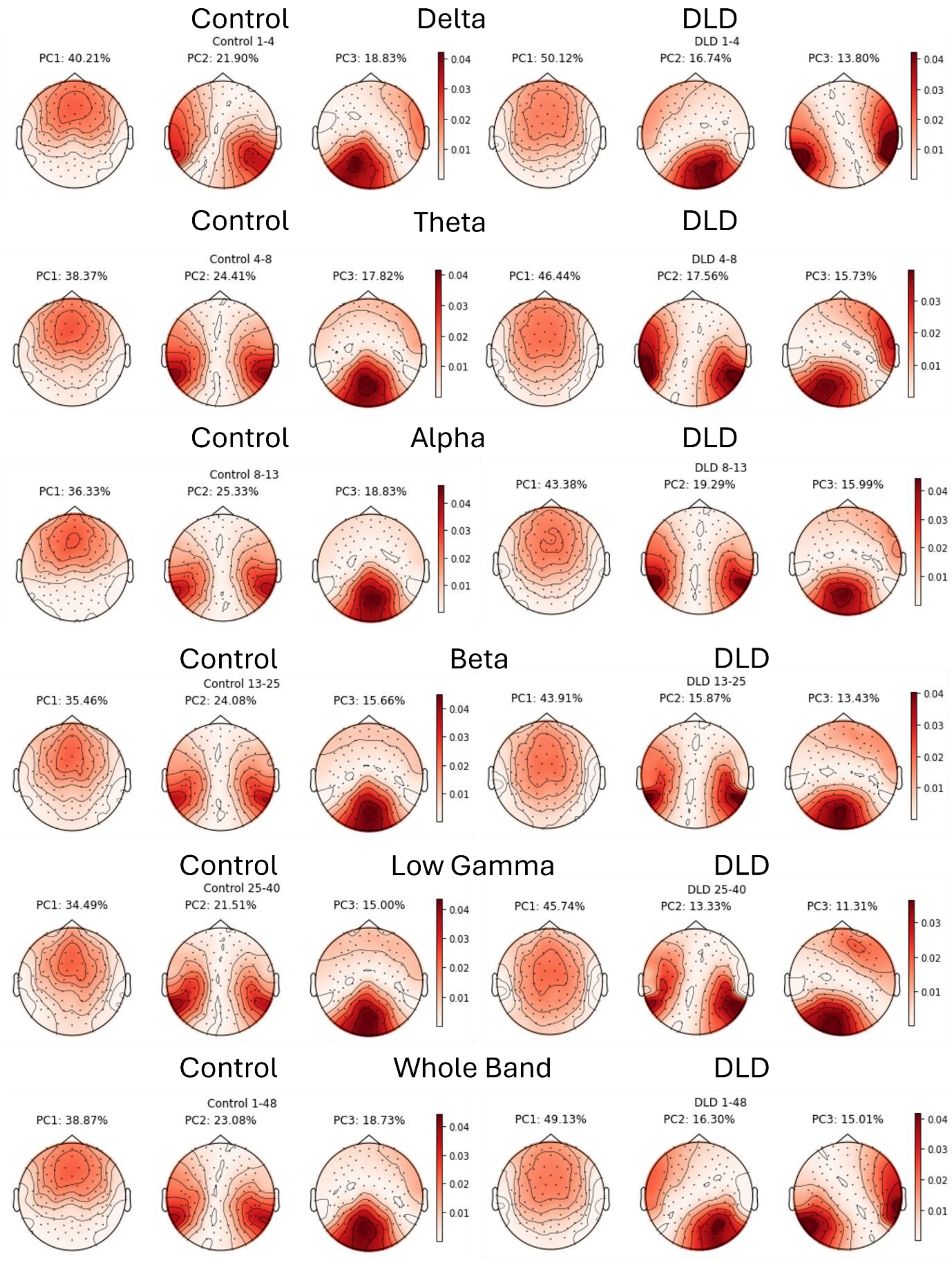

3. ***Table S1.*** *Participant-level Principle Component power ratio for TD and DLD groups*

| Frequency band pair | Data | group | mean | STD | p-value (MWU test / t-test) |
|---|---|---|---|---|---|
| Theta / Delta | PC1 | Control | 0.834 | 0.155 | 0.668(t-test) |
| | | DLD | 0.859 | 0.161 | |
| | PC2 | Control | 0.927 | 0.122 | 0.921(t-test) |
| | | DLD | 0.932 | 0.138 | |
| | PC3 | Control | 0.885 | 0.146 | 0.080(t-test) |
| | | DLD | 0.993 | 0.178 | |
| | EEG | Control | 0.860 | 0.133 | 0.517(t-test) |
| | | DLD | 0.891 | 0.128 | |
| Low Gamma / Delta | PC1 | Control | 0.046 | 0.037 | 0.462(MWU test) |
| | | DLD | 0.034 | 0.022 | |
| | PC2 | Control | 0.071 | 0.049 | 0.158(MWU test) |
| | | DLD | 0.049 | 0.022 | |
| | PC3 | Control | 0.075 | 0.058 | 0.720(MWU test) |
| | | DLD | 0.065 | 0.028 | |
| | EEG | Control | 0.062 | 0.044 | 0.720(MWU test) |
| | | DLD | 0.048 | 0.021 | |
| Low Gamma / Theta | PC1 | Control | 0.053 | 0.041 | 0.396(MWU test) |
| | | DLD | 0.039 | 0.024 | |
| | PC2 | Control | 0.075 | 0.050 | 0.169(MWU test) |
| | | DLD | 0.053 | 0.022 | |
| | PC3 | Control | 0.082 | 0.061 | 0.895(MWU test) |
| | | DLD | 0.066 | 0.027 | |
| | EEG | Control | 0.070 | 0.047 | 0.510(MWU test) |
| | | DLD | 0.053 | 0.023 | |

4. ***Table S2.*** *Shapiro–Wilk Normality Test Results for Participant-Level zMI Distributions*

| Frequency band pair | Data | group | p-value (SW test) |
|---|---|---|---|
| Theta - Delta | PC1 - mean | Control | 0.083 |
| | | DLD | 0.190 |
| | PC1 - var | Control | 0.184 |
| | | DLD | 0.331 |
| | PC2 - mean | Control | 0.502 |
| | | DLD | 0.451 |
| | PC2 - var | Control | 0.519 |
| | | DLD | 0.446 |
| | PC3 - mean | Control | 0.402 |
| | | DLD | **0.032** |
| | PC3 - var | Control | 0.684 |
| | | DLD | 0.550 |
| Low Gamma - Delta | PC1 - mean | Control | 0.073 |
| | | DLD | 0.143 |
| | PC1 - var | Control | **0.012** |
| | | DLD | 0.717 |
| | PC2 - mean | Control | 0.124 |
| | | DLD | **0.013** |
| | PC2 - var | Control | 0.244 |
| | | DLD | 0.124 |
| | PC3 - mean | Control | **0.005** |
| | | DLD | 0.481 |
| | PC3 - var | Control | 0.768 |
| | | DLD | 0.129 |

5. ***Figure S3.*** *Power Spectrum in Filtered Data Across Frequency Bands and filter scalp map*

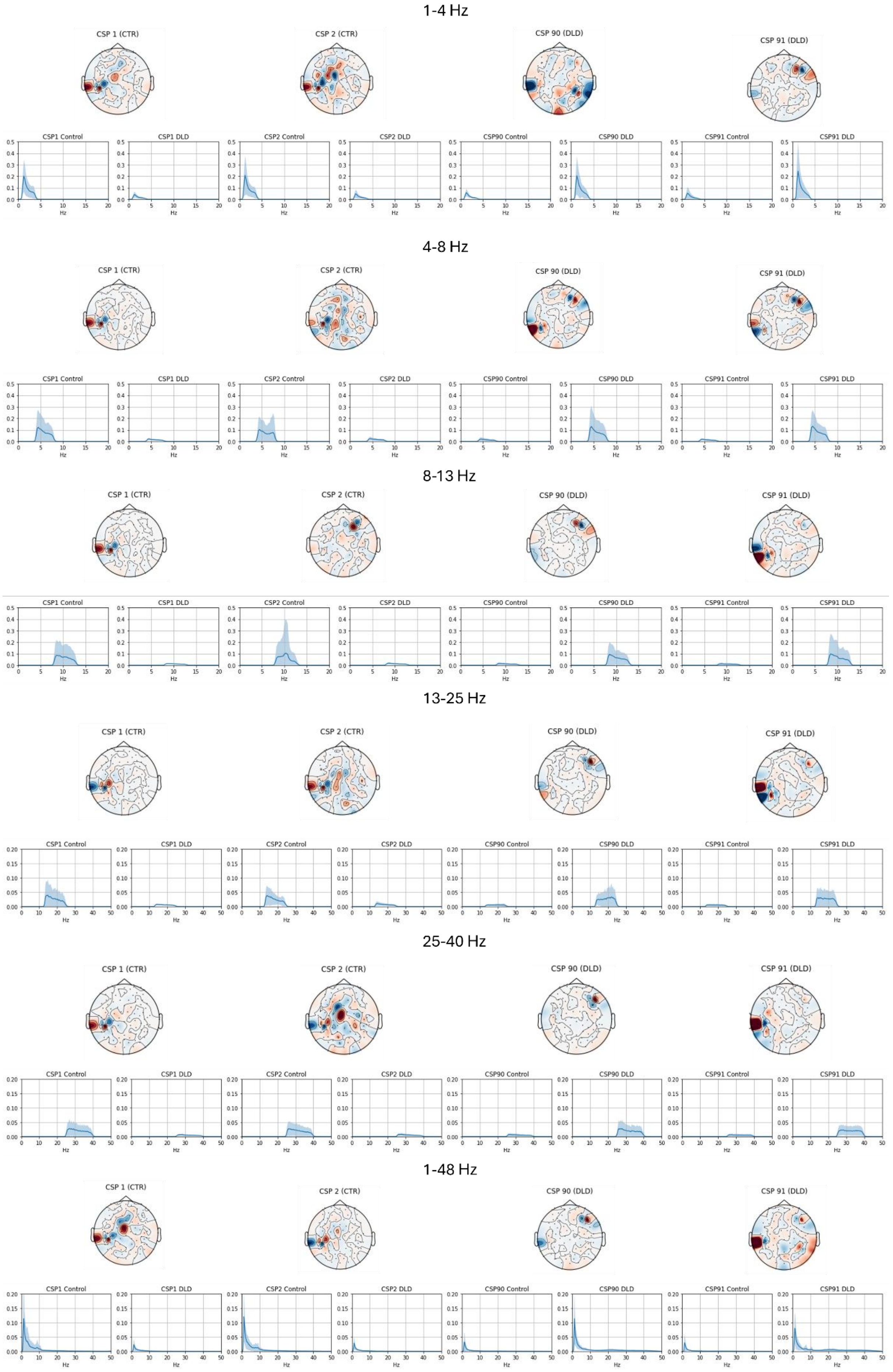

4. ***Figure S4.*** *Complete set of PCA–CSP overlap maps for all frequency bands and groups.*

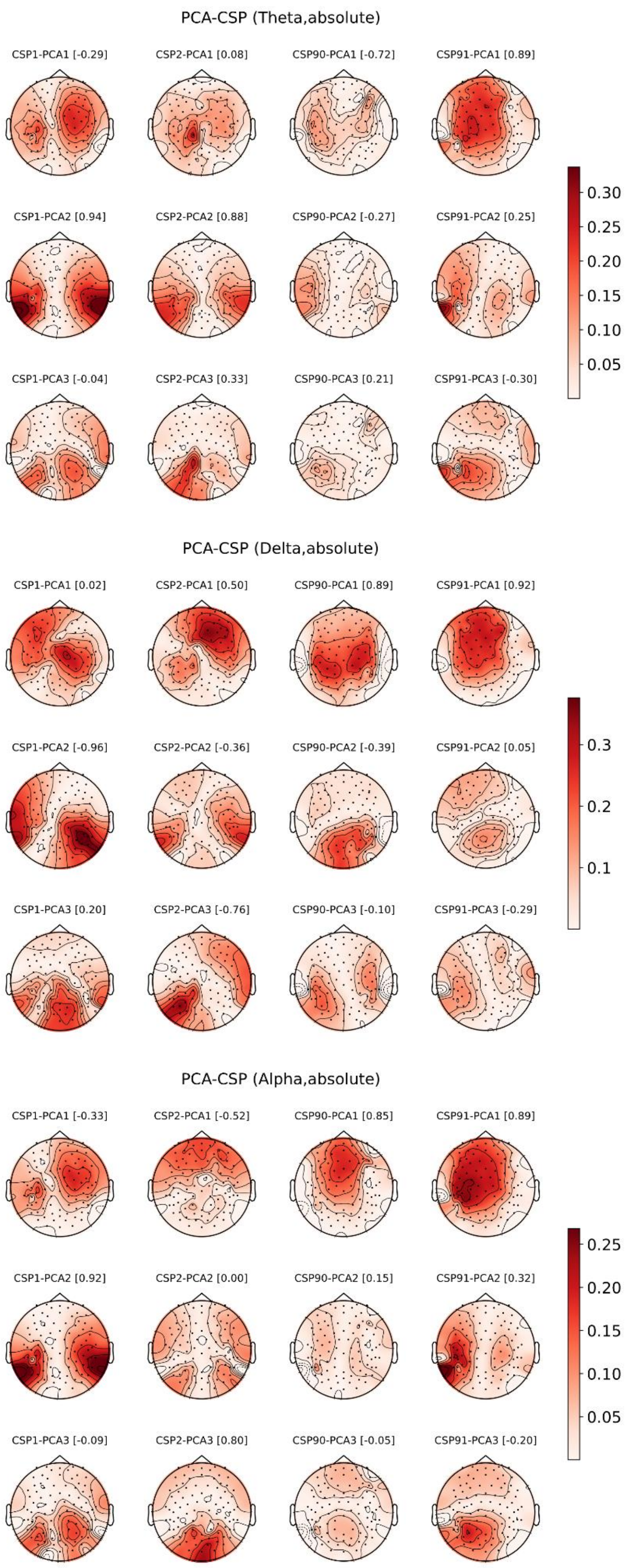

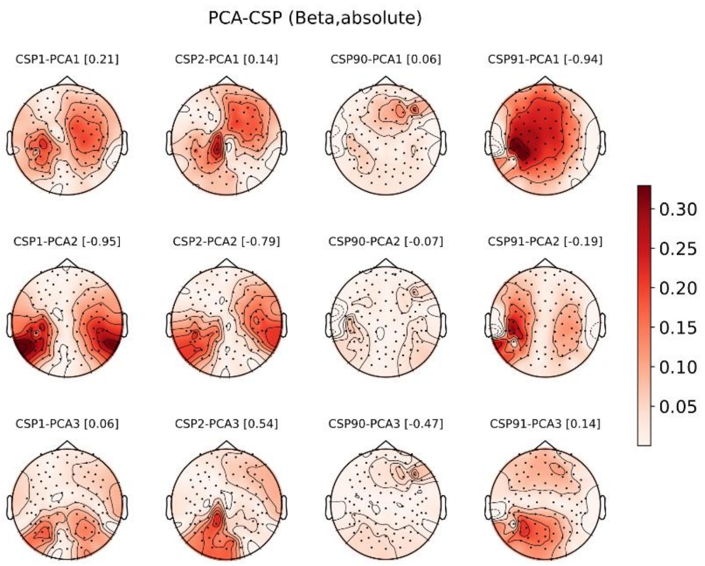
PCA-CSP (Beta,absolute)
CSP1-PCA1 [0.21]
CSP2-PCA1 [0.14]
CSP90-PCA1 [0.06]
CSP91-PCA1 [-0.94]
CSP1-PCA2 [-0.95]
CSP2-PCA2 [-0.79]
CSP90-PCA2 [-0.07]
CSP91-PCA2 [-0.19]
CSP1-PCA3 [0.06]
CSP2-PCA3 [0.54]
CSP90-PCA3 [-0.47]
CSP91-PCA3 [0.14]
0.30
0.25
0.20
0.15
0.10
0.05

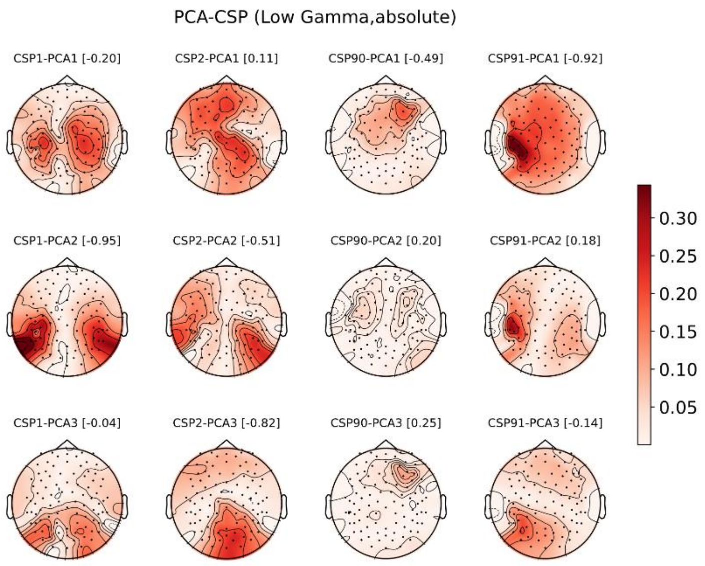
PCA-CSP (Low Gamma,absolute)
CSP1-PCA1 [-0.20]
CSP2-PCA1 [0.11]
CSP90-PCA1 [-0.49]
CSP91-PCA1 [-0.92]
CSP1-PCA2 [-0.95]
CSP2-PCA2 [-0.51]
CSP90-PCA2 [0.20]
CSP91-PCA2 [0.18]
CSP1-PCA3 [-0.04]
CSP2-PCA3 [-0.82]
CSP90-PCA3 [0.25]
CSP91-PCA3 [-0.14]
0.30
0.25
0.20
0.15
0.10
0.05

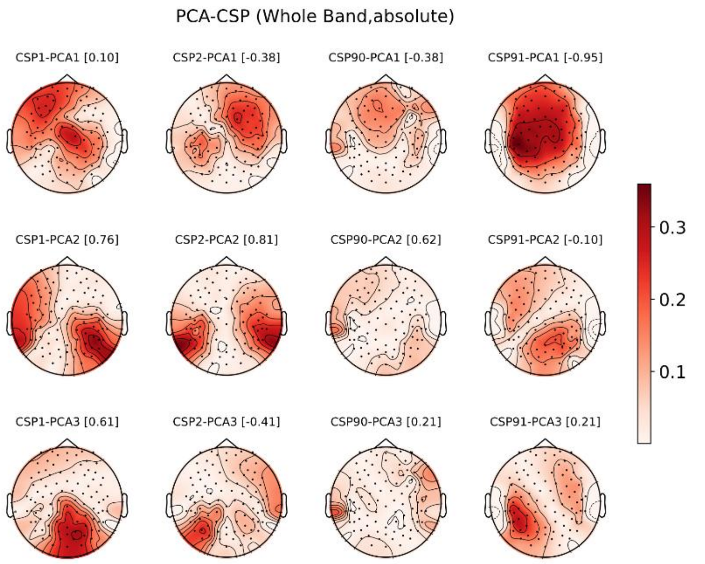
PCA-CSP (Whole Band,absolute)
CSP1-PCA1 [0.10]
CSP2-PCA1 [-0.38]
CSP90-PCA1 [-0.38]
CSP91-PCA1 [-0.95]
CSP1-PCA2 [0.76]
CSP2-PCA2 [0.81]
CSP90-PCA2 [0.62]
CSP91-PCA2 [-0.10]
CSP1-PCA3 [0.61]
CSP2-PCA3 [-0.41]
CSP90-PCA3 [0.21]
CSP91-PCA3 [0.21]
0.3
0.2
0.1
